# Regeneration of dorsal spinal cord neurons after injury via *in situ* NeuroD1-mediated astrocyte-to-neuron conversion

**DOI:** 10.1101/818823

**Authors:** Brendan Puls, Yan Ding, Fengyu Zhang, Mengjie Pan, Zhuofan Lei, Zifei Pei, Mei Jiang, Yuting Bai, Cody Forsyth, Morgan Metzger, Tanvi Rana, Lei Zhang, Xiaoyun Ding, Matthew Keefe, Alice Cai, Austin Redilla, Michael Lai, Kevin He, Hedong Li, Gong Chen

**Affiliations:** All authors part of the Department of Biology, Huck Institutes of Life Sciences, The Pennsylvania State University, University Park, PA 16802, USA

**Author notes:** **Correspondence should be addressed to:** Gong Chen, PhD, Professor and Verne M. Willaman Chair in Life Sciences, Department of Biology, Huck Institutes of Life Sciences, The Pennsylvania State University, University Park, PA 16802, USA, Or Hedong Li, PhD, Department of Biology, Huck Institutes of Life Sciences, The Pennsylvania State University, University Park, PA 16802, USA. Co-first author (Brendan Puls, Yan Ding). Co-senior author (Hedong Li, Gong Chen).

**Keywords:** Spinal cord, NeuroD1, astrocyte, neuronal conversion, in vivo reprogramming

## Abstract

Spinal cord injury (SCI) often leads to impaired motor and sensory functions, partially because the injury-induced neuronal loss cannot be easily replenished through endogenous mechanisms. *In vivo* neuronal reprogramming has emerged as a novel technology to regenerate neurons from endogenous glial cells by forced expression of neurogenic transcription factors. We have previously demonstrated successful astrocyte-to-neuron conversion in mouse brains with injury or Alzheimer’s disease by overexpressing a single neural transcription factor *NeuroD1* via retroviruses. Here we demonstrate regeneration of dorsal spinal cord neurons from reactive astrocytes after SCI via adeno-associated virus (AAV), a more clinically relevant gene delivery system. We find that *NeuroD1* converts reactive astrocytes into neurons in the dorsal horn of stab-injured spinal cord with high efficiency (∼95%). Interestingly, *NeuroD1*-converted neurons in the dorsal horn mostly acquire glutamatergic neuronal subtype, expressing spinal cord-specific markers such as Tlx3 but not brain-specific markers such as Tbr1, suggesting that the astrocytic lineage and local microenvironment affect the cell fate of conversion. Electrophysiological recordings show that the *NeuroD1*-converted neurons can functionally mature and integrate into local spinal cord circuitry by displaying repetitive action potentials and spontaneous synaptic responses. We further show that *NeuroD1*-mediated neuronal conversion can occur in the contusive SCI model, allowing future studies of evaluating this reprogramming technology for functional recovery after SCI. In conclusion, this study may suggest a paradigm shift for spinal cord repair using *in vivo* astrocyte-to-neuron conversion technology to generate functional neurons in the grey matter.

## Introduction

Spinal cord injury (SCI) is a devastating central nervous system (CNS) disorder and often leads to loss of motor and sensory functions below the injury site, even paralysis depending on the severity of the injury (Adams and Hicks, 2005). The pathophysiological process after SCI is rather complex, resulting in neuronal loss, neuroinflammation, demyelination, and Wallerian degeneration of the axons (Norenberg et al., 2004). Reactive astrogliosis is common to CNS injury, and particularly severe after SCI. Resident astrocytes react to injury-induced cytokines and dramatically upregulate the expression of a number of proteins such as the astrocytic marker GFAP and the neural progenitor markers Nestin and Vimentin (Sofroniew, 2009). These reactive astrocytes also become proliferative and hypertrophic in cell morphology. In the acute phase of SCI, reactive astrocytes play important roles in repairing the blood-spinal cord barrier and restricting the size of the primary injury (Herrmann et al., 2008; Okada et al., 2006). However, in the sub-acute or chronic phase, reactive astrocytes constitute the major component of the glial scar, a dense tissue structure that is inhibitory to axonal regeneration (Silver and Miller, 2004). Therefore, for decades, substantial effort has been made to overcome the glial scar and promote regrowth of severed axons through the injury site (Filous and Schwab, 2017). On the other hand, the spinal neurons that are lost during and after the injury need to be replaced in order to rebuild the local neuronal circuits. In this regard, stem cell transplantation has been reported to achieve certain success (Lu et al., 2017; Tuszynski et al., 2014), however long-term survival of transplanted cells in the injury site has been an unsolved issue (Goldman, 2016).

*In vivo* neuronal reprogramming has recently emerged as a novel technology to regenerate functional new neurons from endogenous glial cells by overexpressing neurogenic transcription factors in the CNS (Barker et al., 2018; Gascon et al., 2016; Grande et al., 2013; Guo et al., 2014; Lei et al., 2019; Li and Chen, 2016; Liu et al., 2015; Niu et al., 2013). In the injured spinal cord, a combination of growth factor treatment and forced expression of the neurogenic transcription factor *Neurogenin 2* (*Ngn2*) has been reported to stimulate neurogenesis from neural progenitors (Ohori et al., 2006). However, these newly generated neurons suffer from poor long-term survival. More recently, the transcription factor *Sox2* has been shown to reprogram astrocytes into proliferating neuroblasts, which require further treatment with a histone deacetylase inhibitor, valproic acid (VPA), to differentiate into mature neurons (Su et al., 2014). With additional treatment of neurotrophic factors, *Sox2*-converted neurons can express several neuronal subtype markers, but predominately VGluT, a marker for glutamatergic neurons (Wang et al., 2016).

Our group has previously demonstrated that the neurogenic transcription factor *NeuroD1* can reprogram reactive astrocytes into functional neurons in the stab-injured brain and in a mouse model for Alzheimer’s disease (Guo et al., 2014). Following studies demonstrated that NeuroD1-mediated *in vivo* astrocyte-to-neuron conversion can reverse glial scar back to neural tissue (Zhang et al., 2018) and repair the damaged motor cortex after ischemic stroke (Chen et al., 2018). The major goal of the current study is to determine whether *NeuroD1* can reprogram reactive astrocytes into functional neurons in the injured spinal cord. Using adeno-associated virus (AAV) for gene delivery and a *Cre-Flex* system with *GFAP* promoter to target reactive astrocytes specifically, our results indicate that *NeuroD1* can mediate direct astrocyte-to-neuron conversion with high efficiency in both stab and contusive SCI models. The *NeuroD1*-converted neurons preferentially acquire glutamatergic phenotype in the dorsal horn and express neuronal subtype markers specific to the spinal cord such as Tlx3. Patch clamp recordings further demonstrate that the *NeuroD1*-converted neurons can functionally mature and integrate into spinal cord circuitry. Together, our results indicate that *NeuroD1*-mediated neuronal conversion opens an avenue to treat SCI with internal glial cells through AAV-based gene therapy that may regenerate a diversity of neuronal subtypes for functional repair.

## Materials and methods

### Animal use

GAD-GFP mice (Tg[Gad1-EGFP]94Agmo/J) and wild-type C57BL/6 mice were purchased from the Jackson Laboratory and bred in house. Mice of 2-4 months old (both male and female) were used. Mice were housed in a 12 hr light/dark cycle and supplied with sufficient food and water. All animal use and studies were approved by the Institutional Animal Care and Use Committee (IACUC) of the Pennsylvania State University. All procedures were carried out in accordance with the approved protocols and guidelines of National Institute of Health (NIH).

### Retrovirus and AAV production

Retroviral vectors expressing *GFP* and *NeuroD1-GFP* under the *CAG* promoter (*pCAG*) were previously described (Guo et al., 2014). Retrovirus packaging, purification and titering were performed as previously described (Guo et al., 2014).

For AAV-mediated gene expression, the *Cre-Flex* system was applied to target transgene expression specifically to reactive astrocytes using the *human GFAP* (*hGFAP*) promoter. To generate *pAAV-hGFAP∷Cre* vector, the *hGFAP* promoter was first amplified from *pDRIVE-hGFAP* plasmid (InvivoGen) by PCR and inserted into *pAAV-MCS* (Cell Biolab) between the MluI and SacII sites to replace the *CMV* promoter. The *Cre* gene coding fragment was then similarly subcloned from *phGFAP-Cre* (Addgene plasmid #40591) and inserted into *pAAV MCS* between the EcoRI and SalI sites. To construct *pAAV-FLEX* vectors expressing transgenes, the coding sequences of *NeuroD1, mCherry* or *GFP* were amplified by PCR from the corresponding retroviral constructs. The *NeuroD1* fragment was fused with either *P2A-mCherry* or *P2A-GFP* and subcloned into the *pAAV-FLEX-GFP* vector (Addgene plasmid #28304) between the KpnI and XhoI sites. All plasmid constructs were confirmed by sequencing. The AAV-CamKII-GFP plasmid was purchased from Addgene (#64545). For AAV production, HEK 293T cells were transfected with the *pAAV* expression vectors, *pAAV9-RC* vector (Cell Biolab), and *pHelper* vector (Cell Biolab) to generate AAV particles carrying our transgenes. Three days after transfection, the cells were scraped in their medium and centrifuged. The supernatant was then discarded and the cell pellet was frozen and thawed four times, resuspended in a discontinuous iodixanol gradient, and centrifuged at 54,000 rpm for two hours. Finally, the virus-containing layer was extracted, and the viruses were concentrated using Millipore Amicon Ultra Centrifugal Filters. The viral titers were determined using the QuickTiter™ AAV Quantitation Kit (Cell Biolabs) and then diluted to a final concentration of 1×10^10^ genome copy (GC)/ml for injection.

### Laminectomy, injury, and stereotaxic viral injection

Mice were anesthetized by intraperitoneal injection of ketamine/xylazine (80-120 mg/kg ketamine; 10-16 mg/kg xylazine). A laminectomy was then performed at the T11-T12 vertebrae to expose dorsal surface of the spinal cord, and a stab or contusion injury was performed. The stab injury was conducted with a 31-gauge needle at the center of the exposed surface, 0.4 mm lateral to the central artery with a depth of 0.4 mm, while the contusion injury was generated with a force of 45 kdyn on an Infinite Horizon Impactor (IH-0400, Precision Systems and Instrumentation) directly at the center of the exposed surface. Either immediately following the injury or at a specified delay, 1.0 μL of concentrated virus was injected using a 50 μl Hamilton syringe with a 34-gauge injection needle at a rate of 0.05 μL/min at the same coordinates for the stab injury or 1 mm away for the contusion injury. The needle was then kept in place for three minutes after injection to prevent drawing out the virus during withdrawal. The surgical area was then treated with antibiotic ointment and the skin was clipped for a week to allow the skin to re-suture. The mice were kept on a heating pad and treated with Carprofen for pain relieve via subcutaneous injection (5 mg/kg) on the day of surgery and drinking water (10 mg/kg) for three days after surgery and closely monitored for one week to ensure full recovery of health.

### Electrophysiology

Mice were sacrificed at defined time points by anesthetization with 2.5% Avertin and decapitation. The spinal cord segment was then removed from the spine into cutting solution (125 mM NaCl, 2.5 mM KCl, 1.3 mM MgSO_4_, 26 mM NaHCO_3_, 1.25 mM NaH_2_PO_4_, 2.0 mM CaCl_2_ and 10 mM glucose adjusted to pH 7.4 and 295 mOsm/L and bubbled for one hour with 95% O_2_/5% CO_2_) cooled on ice, where it was encased in an agarose matrix (Sigma) and cut into 300 µm thickness slices using a VT3000 vibratome (Leica). Slices were then incubated for one hour in holding solution (92 mM NaCl, 2.5 mM KCl, 1.25 mM NaH_2_PO_4_, 30 mM NaHCO_3_, 20 mM HEPES, 15 mM glucose, 12 mM N-Acetyl-L-cysteine, 5 mM Sodium ascorbate, 2 mM Thiourea, 3 mM Sodium pyruvate, 2 mM MgSO_4_, and 2 mM CaCl_2_, adjusted to pH 7.4 and 295 mOsm/L and bubbled continuously with 95% O_2_/5% CO_2_) at room temperature before patch-clamp recording in standard ACSF (125 mM NaCl, 2.5 mM KCl, 1.25 mM NaH_2_PO_4_, 26 mM NaHCO_3_, 1.3 MgSO_4_, 2.5 mM CaCl_2_, and 10 mM glucose adjusted to pH 7.4 and 295 mOsm/L and bubbled for one hour with 95% O_2_/5% CO_2_). Both native and converted cells were recorded by whole-cell recording using standard inner solution (135 mM K-gluconate, 10 mM KCl, 5 mM Na-phosphocreatine, 10 mM HEPES, 2 mM EGTA, 4 mM MgATP, and 0.5 mM Na_2_GTP, adjusted to pH 7.4 and 295 mOsm/L) with the membrane potential held at −70 mV. Typical values for the pipette and total series resistances were 2-10 MΩ and 20-60 MΩ, respectively. Data were collected using the pClamp 9 software (Molecular Devices) by sampling at 10 kHz and filtering at 1 kHz. Data were then analyzed and plotted with the Clampfit 9.0 software (Molecular Devices).

### Immunohistochemistry, immunocytochemistry, and microscopy

After perfusion, the target region of the spinal cord (∼ 0.5 cm in length) was surgically dissected, fixed in 4% paraformaldehyde (PFA) in PBS for one day, dehydrated in 30% sucrose solution for one day, and sectioned into 30 μm coronal or horizontal slices using a Leica CM1950 cryostat. The slices were collected serially in 24-well plates so that distance from the injury site could later be ascertained. The samples were then stored at 4 °C in 0.02% sodium azide (NaN_3_) in PBS to prevent bacterial degradation. Spinal cord slices were chosen for immunohistochemistry based on infection of the dorsal horn by inspecting the reporter protein (mCherry or GFP) in the storage solution under a fluorescent microscope. For the stab injury experiments, care was taken to select coronal slices at least 100 μm from the injury site to ensure tissue integrity. On the first day of staining, samples were washed in PBS three times for five minutes per wash, permeablized with 2% Triton X-100 in PBS for 20 minutes, and blocked using a 5% normal donkey serum (NDS) and 0.1% Triton-X in PBS for two hours to reduce non-specific binding of the antibodies. The samples were then incubated with primary antibodies diluted in the same blocking buffer at 4 °C for two nights to allow thorough penetration of the antibodies. On the third day, the samples were recovered to room temperature, washed in PBS three times for five minutes per wash, and incubated with secondary antibodies diluted in blocking buffer for one hour. Finally, the samples were washed in PBS three more times for ten minutes per wash and mounted on glass slides with coverslips using anti-fading mounting solution (Invitrogen). The immunostained samples were examined and imaged using Olympus FV1200 and Zeiss LSM 800 laser confocal microscopes. Z-stacks were collected for the *in vivo* images for the whole thickness of the samples and maximum intensity and z-stack projections were used for image preparation and analysis.

### Quantification and data analysis

As a result of our carefully selected injection coordinates described above, infected cells were mostly found in the dorsal horn of the spinal cord, Rexed laminae 1-6 (Rexed, 1954). For most of the quantification, including cell conversion and NeuN acquisition, cells were counted if they appeared in any part of this region. For quantification based on cell subtype (Figs. 3, 4), cells were only counted if they appeared in Rexed laminae 1-3, centered about the substantia gelatinosa, a region dominated by small, excitatory interneurons and easily demarcated due to its high cell density (Santos et al., 2007). This region was chosen for its ease of demarcation and so that a consistent local population of neuronal subtypes could be expected from sample to sample. Quantification was performed on collected images using the z-stacked images as a guide and the layered stacks to check the vertical dimension. Strict background cutoffs for positive signals were calculated for each channel as three times the average background intensity for the relevant tissue and antibody. Cells were binned by presence (i.e., above the background cutoff) or absence (i.e., below the background cutoff) for each marker in question, using the viral fluorophore (mCherry or GFP) to identify infected cells and DAPI to confirm each cell for counting. To estimate the total number of converted neurons per infection for our contusion experiments, we multiplied the average number of NeuN+ infected cells per horizontal section, calculated from one dorsal, one central, and one ventral section, by the total number of horizontal sections per sample. All quantification was performed on three biological replicates per data point and is reported as the means and standard deviations of the three replicates.

**Figure 1:**
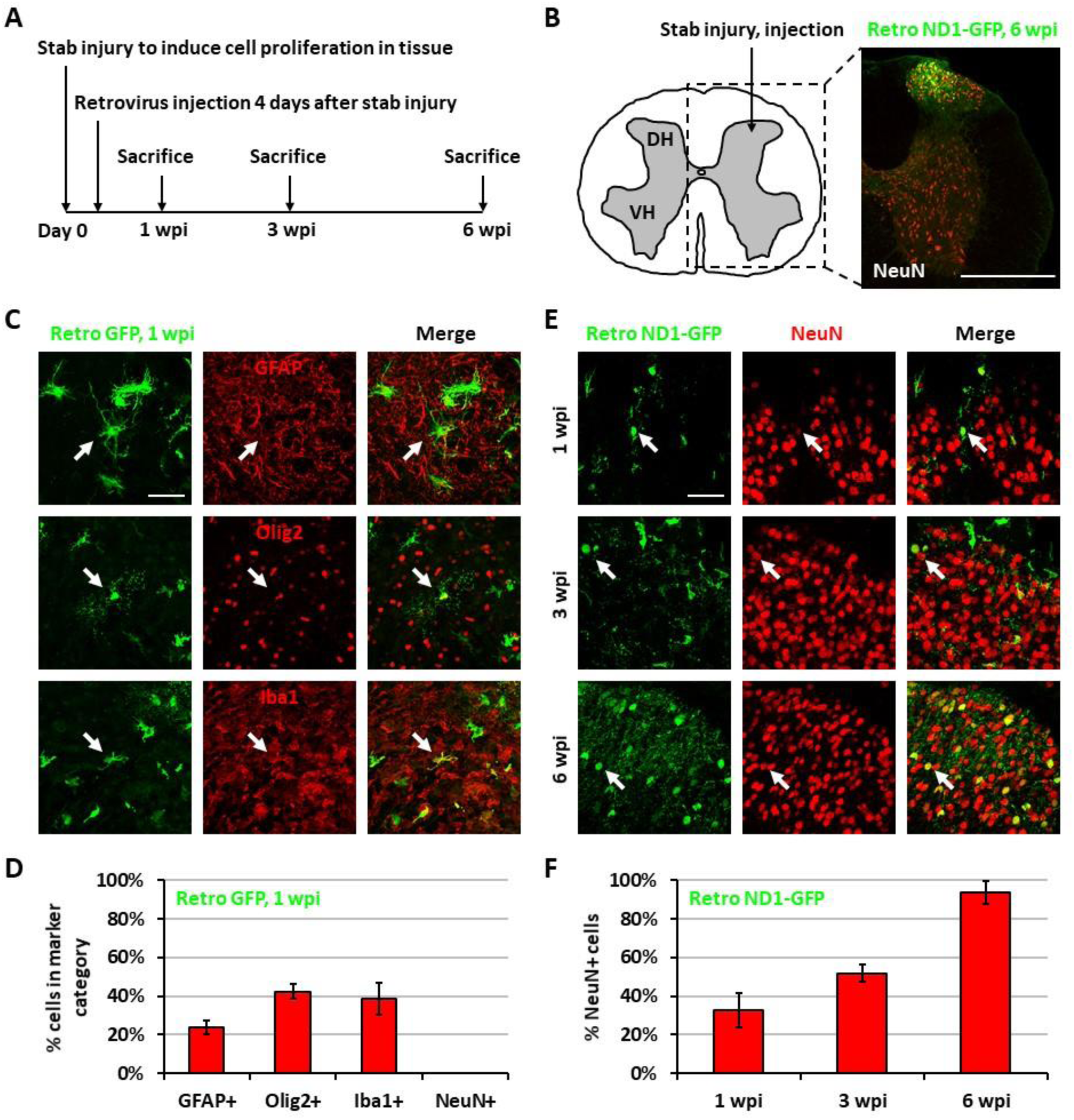
Neuronal conversion using NeuroD1-expressing retrovirus after stab injury in the spinal cord dorsal horn. (**A**) Experiment paradigm. (**B**) Schematic of dorsal horn injury and injection. Coordinates are 0.4 mm lateral of central artery and 0.4 mm below the tissue surface. Stab injury was performed with a 32-guage needle followed by stereotaxic injection at the injury site. Scale bar 500 µm. (**C**) Three main types of proliferating cells at 1 wpi after injection of Retro GFP: astrocytes, OPCs, and microglia. GFAP, Olig2, and Iba1 staining show markers for these cell types. Arrows show examples of each. Scale bar 50 µm. (**D**) Quantification based on staining for Retro GFP, 1 wpi samples. Bars show the mean and standard deviation of three replicates. (**E**) Converting cells in the dorsal horn 1 wpi, 3 wpi, and 6 wpi after Retro ND1-GFP injection. Maturing cells gradually up-regulate NeuN, adapt neuronal morphology, and reorganize. Arrows show example NeuN+ cells. Scale bar 500 µm. (**F**) Quantification based on staining for Retro ND1-GFP, 1W samples. Bars show the mean and standard deviation of three replicates.

**Figure 2:**
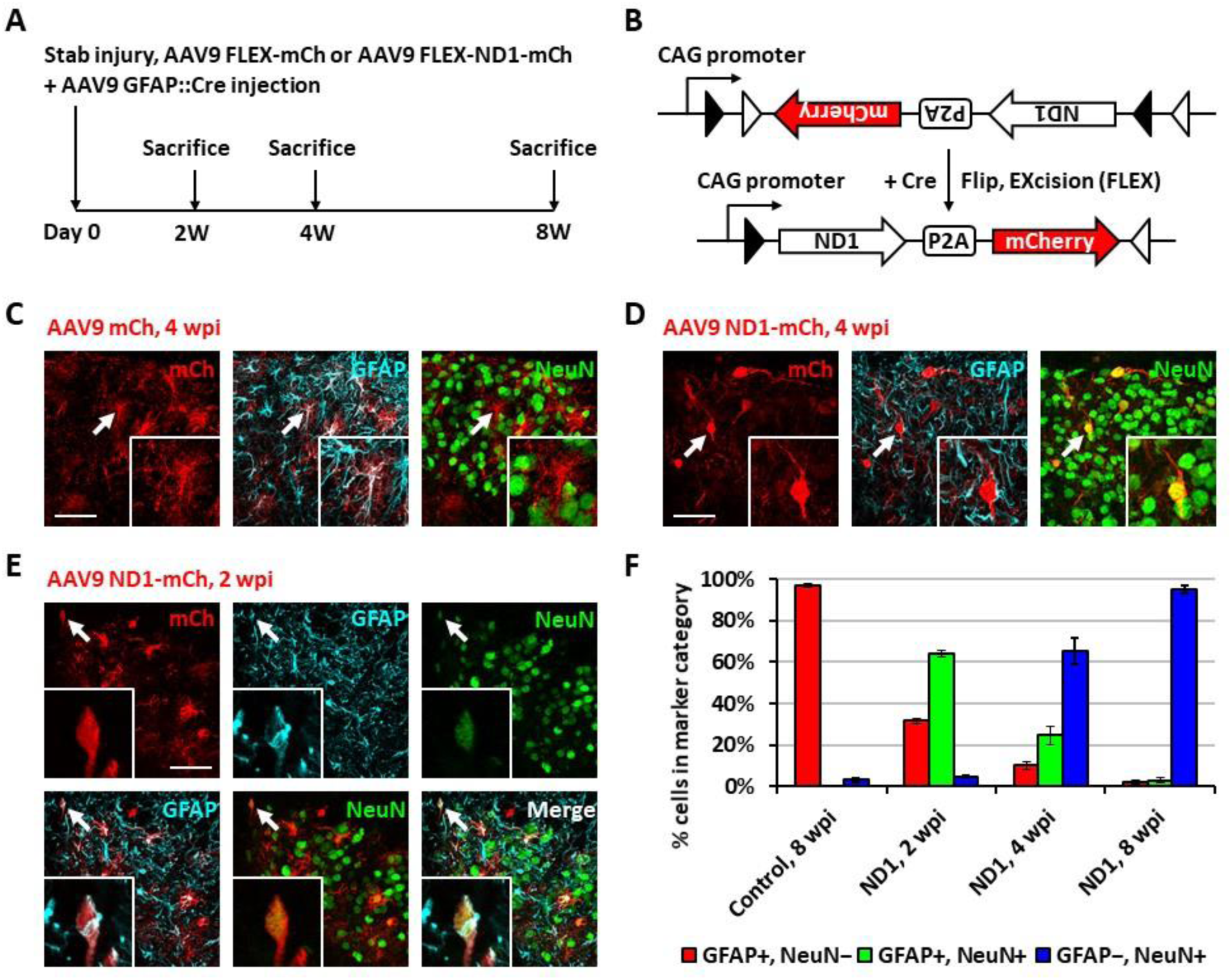
Neuronal conversion using NeuroD1-expressing AAV9 after stab injury in the spinal cord dorsal horn. (**A**) Experiment paradigm. (**B**) AAV9 FLEX-NeuroD1-mCherry/AAV9 GFAP∷Cre system (abbreviated elsewhere as AAV9 ND1-mCh). GFAP promoter restricts infected cells to astrocytes. Control virus replaces the ND1 transgene with an additional mCh. (**C**) Infected astrocytes in the dorsal horn 4 wpi after AAV9 mCh injection. Arrows and inset (2×mag) show an example GFAP+ cell. Scale bar 50 µm. (**D**) Converted neurons in the dorsal horn 4 wpi after AAV9-ND1-mChy injection. Arrows and inset (2×mag) show an example NeuN+ cell. Scale bar 50 µm. (**E**) Converting cells in the dorsal horn 2 wpi after AAV9-ND1-mChy injection. Arrows and inset (4×mag) show an example NeuN+/GFAP+ cell. Scale bar 50 µm. (**F**) Quantification based on staining for AAV9 ND1-mCh, 2 wpi and 4 wpi, and AAV9 ND1-GFP or AAV9 GFP (Control), 8 wpi samples. Bars show the mean and standard deviation of three replicates. Infected cells at 2 wpi are mostly transitional, staining positive for NeuN and GFAP, and by 4 wpi are mostly converted, staining positive for only NeuN.

**Figure 3:**
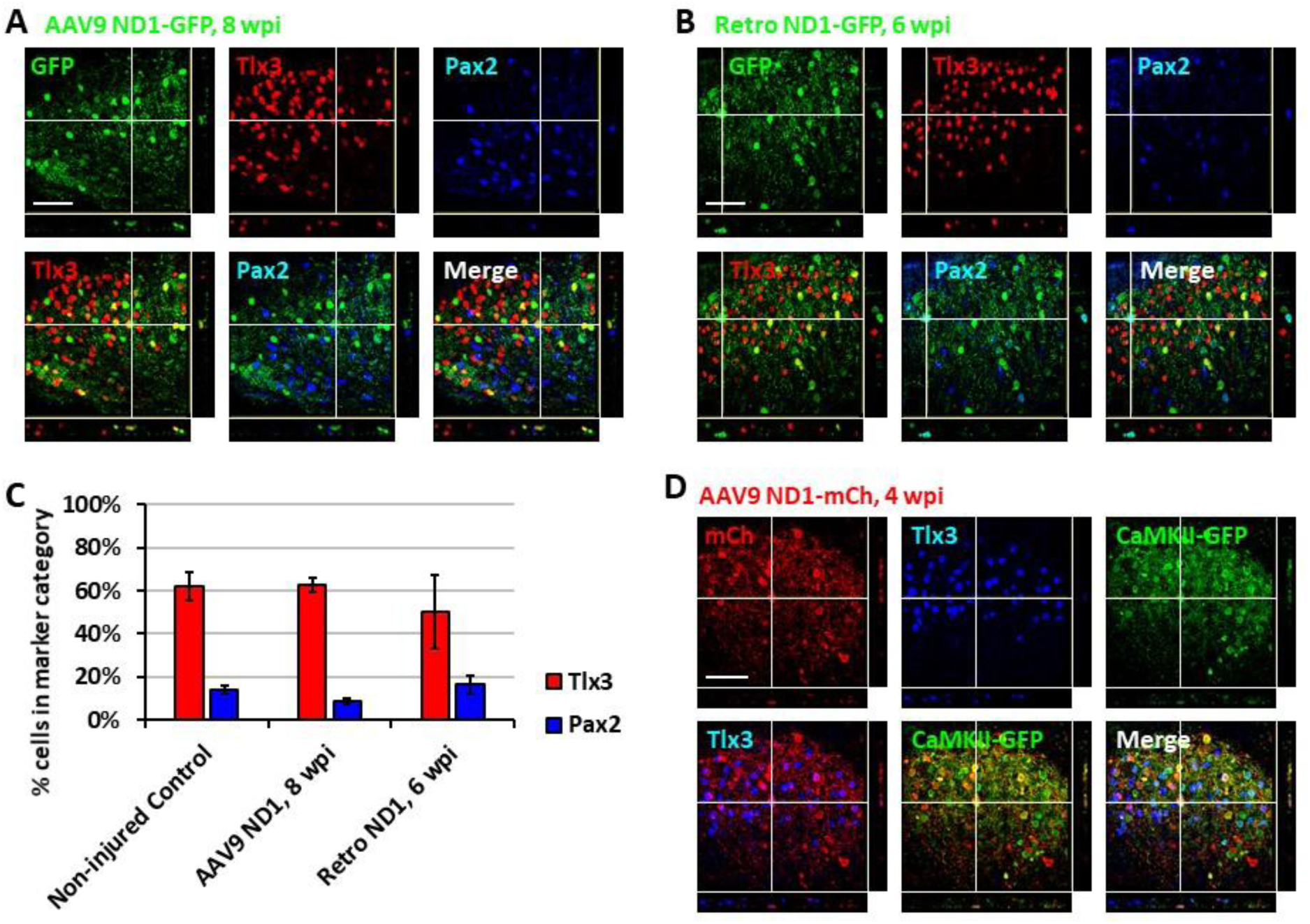
Subtypes of NeuroD1-converted neurons in the spinal cord dorsal horn. (**A**) Tlx3 (glutamatergic) and Pax2 (GABAergic) subtype staining for converted neurons in the dorsal horn 8 wpi after AAV9 ND1-GFP injection. Z-projection targets an example Tlx3+ neuron. Scale bar 50 µm. (**B**) Tlx3 and Pax2 subtype staining for converted neurons in the dorsal horn 6 wpi after Retro ND1-GFP injection. Z-projection targets an example Pax2+ neuron. Scale bar 50 µm. (**C**) Quantification based on subtype staining for AAV9 ND1-GFP, 8 wpi and Retro ND1-GFP, 6 wpi samples. Control data is based on NeuN+ cells in uninjured, untreated tissue. Bars show the mean and standard deviation of three replicates. (**D**) AAV9 ND1-mCy and CaMK2-GFP co-injection at 4 wpi with strong (89.5±5.2%) co-labeling of CaMK2 for converted, Tlx3+ cells. Z-projection targets an example Tlx3+, CaMK2+ neuron. Scale bar 50 µm.

**Figure 4:**
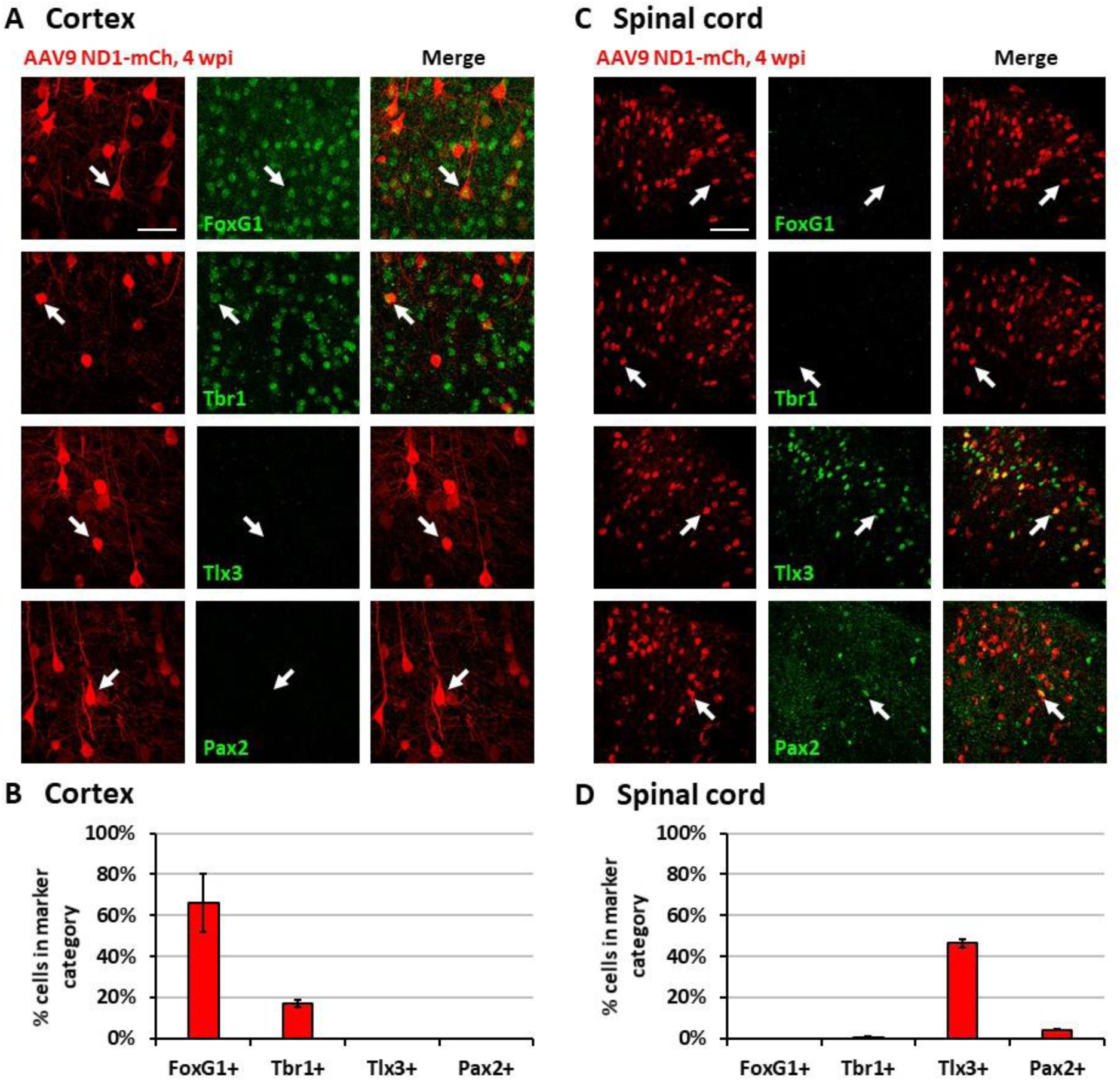
Region-specific subtypes of NeuroD1-converted neurons in the brain versus the spinal cord. (**A**) Subtype staining for converted neurons in the cortex 4 wpi after AAV9 ND1-mCh injection. Arrows show examples of cells positive for each subtype. Scale bar 50 µm. (**B**) Quantification based on subtype staining in the cortex for AAV9 ND1-mCh, 4 wpi samples. Bars show the mean and standard deviation of three replicates. (**C**) Subtype staining for converted neurons in the spinal cord dorsal horn 4 wpi after AAV9 ND1-mChy injection. Arrows show examples of cells positive for each subtype. Scale bar 50 µm. (**D**) Quantification based on subtype staining in the spinal cord for AAV9 ND1-mCh, 4 wpi samples. Bars show the mean and standard deviation of three replicates.

## Results

### NeuroD1 reprograms reactive astrocytes into neurons in the injured spinal cord

SCI has been studied for decades but so far there is still limited therapy to treat SCI patients. Besides axonal degeneration, neuronal loss following SCI is a major obstacle for functional recovery. We previously demonstrated that expressing NeuroD1 in reactive astrocytes after brain injury can directly convert astrocytes into neurons (Guo et al., 2014). In this study, we investigated whether such *in vivo* direct conversion technology can regenerate functional new neurons in injured spinal cord. To target the dividing reactive astrocytes after injury, we employed retroviruses that mainly express ectopic genes in dividing cells but not in neurons, which cannot divide. We injected NeuroD1-expressing retroviruses at 4 days post-stab injury (dpi), when many dividing reactive astrocytes have been detected (Chen et al., 2008; Hong et al., 2014), and analyzed samples at 1, 3, and 6 weeks post-injection (wpi) (Fig. 1A). In this study, we chose the spinal cord dorsal horn as our major region of interest because it is composed of both excitatory and inhibitory neurons and is critical to afferent sensory information processing (Fig. 1B). We are currently investigating motor neuron regeneration in the spinal cord ventral horn in a separate study. We first explored the cell types infected by our control CAG∷GFP retroviruses. At 1 wpi, we found that the control GFP retroviruses infected a mixture of glial cell types including reactive astrocytes (GFAP+ and some GFAP+/Olig2+), oligodendrocyte progenitor cells (OPCs) (Olig2+) and microglia (Iba1+) (Fig. 1C, D), but not NeuN+ neurons (Fig. 1D). In contrast, cells infected by the CAG∷NeuroD1-GFP retrovirus showed an increasing number of NeuN+ cells with neuronal morphology over time (Fig. 1E), and quantitatively reached 93.5% at 6 wpi (Fig. 1F), indicating a successful glia-to-neuron conversion in the injured spinal cord.

While retroviruses can target reactive glial cells, the number of dividing glial cells at the time of viral infection may be limited. In order to move our *in vivo* reprogramming technology towards clinical settings in the future, we adopted an AAV gene delivery system in which the transgene expression is controlled by an astrocyte-specific *GFAP* promoter (Fig. 2A). Specifically, we used a *Cre-Flex* gene expression system, which contains two AAV vectors, with one encoding *GFAP-Cre* and the other encoding the transgene in reverse form flanked by double LoxP sites (FLEX vector) (Atasoy et al., 2008; Chen et al., 2018; Liu et al., 2015). Thus, when the two AAVs are co-injected into the spinal cord, Cre recombinase will be expressed in the infected reactive astrocytes and turn on the transgene expression in FLEX vector by flipping the transgene sequence into the correct form for transcription (Fig. 2B). We first confirmed the specificity of the *Cre-Flex* AAV system in the spinal cord by co-injecting AAV GFAP∷Cre and AAV FLEX-CAG∷mCherry (or ∷GFP) into the stab-injured dorsal horn. The control virus infected cells were mostly GFAP+, NeuN− astrocytes at 4 wpi (Fig. 2C). Next, we co-injected AAV GFAP∷Cre with AAV FLEX-CAG∷NeuroD1-P2A-mCherry into the stab-injured dorsal horn. In contrast to the control AAV, the NeuroD1-mCherry infected cells were mostly NeuN+/GFAP− neurons with clear neuronal morphology at 4 wpi (Fig. 2D). NeuroD1 overexpression in the infected cells was confirmed by immunostaining (Supp. Fig. 1). Interestingly, besides NeuN+/GFAP− converted neurons, we also observed many NeuroD1-AAV-infected cells at 2 wpi with co-immunostaining of both GFAP+ and NeuN+ (Fig. 2E), suggesting a potential intermediate stage during astrocyte-to-neuron conversion. We termed these GFAP+/NeuN+ cells induced by NeuroD1 expression in astrocytes as “AtN transitional cells”. We did not observe any such transitional cells in the control mCherry-infected spinal cord after injury, suggesting that AtN conversion does not happen following neural injury but can be induced by ectopic expression of transcription factors such as NeuroD1. Quantitative analysis revealed that the control AAV-infected cells were mostly GFAP+ astrocytes by 8 wpi (Fig. 2F, left red bar), whereas NeuroD1 AAV-infected cells showed a progressive increase in the percentage of neurons (NeuN+/GFAP−, blue bar in Fig. 2F) from 2 to 8 wpi, reaching ∼95% at 8 wpi (Fig. 2F, right blue bar). Note that at 2 wpi, over 60% of NeuroD1-infected cells were GFAP+/NeuN+ transitional cells (green bar in Fig. 2F), which gradually decreased at 4 wpi and 8 wpi together with a decrease of GFAP+ astrocytes (red bar in Fig. 2F) among the NeuroD1-infected cell population. Further analysis showed that neither transitional cells nor converted neurons exhibited significant cell death suggesting that apoptosis does not play a significant role during the NeuroD1-mediated cell conversion process (Supp. Fig. 2). Comparing to Ngn2-mediated or Ascl1-mediated AtN conversion (Gascon et al., 2016), less apoptosis was detected during NeuroD1-mediated conversion process, which may suggest that different transcription factors act through different signaling and metabolic pathways to carry out cell conversion.

### NeuroD1 converts dorsal spinal astrocytes into Tlx3+ glutamatergic neurons

After demonstrating astrocyte-to-neuron conversion in the spinal cord, we next investigated which subtypes of neurons were generated through NeuroD1-mediated conversion. The dorsal horn of the spinal cord contains two main neuronal subtypes: glutamatergic and GABAergic neurons (Abraira and Ginty, 2013). During spinal cord development, two transcription factors, Tlx3 and Pax2, appear to play critical roles in determining cell fate specification in the dorsal horn (Cheng et al., 2005; Huang et al., 2008). Interestingly, by examining AAV NeuroD1-GFP-infected cells in the dorsal horn at 8 wpi, we found that the majority of NeuroD1-converted neurons were Tlx3+ (62.6 ± 3.3%), suggesting a majority glutamatergic neuronal subtype (Fig. 3A). In contrast, only a small percentage of NeuroD1-converted neurons in the dorsal horn were Pax2+ (8.8 ± 1.3%), suggesting a minority GABAergic neuronal subtype (Fig. 3A). Because AAV might infect a small proportion of neurons (Fig. 2F, control), we further examined retrovirus NeuroD1-GFP-infected cells in the dorsal horn at 6 wpi and found that, similar to our AAV experiments, the retrovirus NeuroD1-converted neurons were mainly Tlx3+ (Fig. 3B). As a control, we quantified the proportion of Tlx3+ and Pax2+ neurons in the spinal cord dorsal horn of uninjured and untreated mice, and found that the native proportions of Tlx3+ and Pax2+ cells were 62.0 ± 6.4% and 14.2 ± 2.0%, respectively (Fig. 3C, left two bars). In comparison, the proportion of Tlx3+ and Pax2+ neurons among the AAV NeuroD1-converted neurons were 62.6 ± 3.3% and 8.8 ± 1.3%, respectively (Fig. 3C, middle two bars); and among the retrovirus NeuroD1-converted neurons were 50.3 ± 17.0% of Tlx3+ and 16.4 ± 4.3% Pax2+ (Fig. 3C, right two bars). These results suggest that the majority of NeuroD1-converted neurons in the dorsal horn of spinal cord are Tlx3+ neurons, with a small proportion being Pax2+ neurons.

We further confirmed the neuronal subtypes after NeuroD1 conversion using AAV CaMKII-GFP to identify glutamatergic neurons and GAD-GFP transgenic mice to identify GABAergic neurons. When co-injecting AAV GFAP∷Cre and Flex-NeuroD1-mCherry together with AAV CaMKII∷GFP (Dittgen et al., 2004), we observed 89.5 ± 5.2% (n=3) of GFP+ cells co-expressing Tlx3+, confirming that these Tlx3+ neurons are indeed glutamatergic (Fig. 3D). Many NeuroD1-mCherry converted neurons were also colocalizing with CaMKII-GFP (Fig. 3D), suggesting that they were glutamatergic neurons. When injecting AAV GFAP∷Cre and Flex-NeuroD1-mCherry in GAD-GFP mice (n = 3), in which GABAergic neurons are genetically labeled with GFP, we did not observe GAD-GFP co-expression with Tlx3+, as expected. Indeed, the CaMKII and GAD markers co-stained consistently with endogenous Tlx3+ and Pax2+ neurons as well, which we observed in the dorsal horn of spinal cord in uninjured, untreated mice (Supp. Fig. 3). Therefore, the majority of NeuroD1-converted neurons in the dorsal horn of the spinal cord are glutamatergic neurons, consistent with our finding in the mouse cortex (Chen et al., 2018).

### NeuroD1-converted neurons express region-specific neuronal subtype markers

While NeuroD1-converted neurons appear to be mainly glutamatergic neurons in both the mouse cortex and spinal cord, we further investigated whether they are the same type of glutamatergic neurons or not. For this purpose, we injected the same AAV GFAP∷Cre and AAV FLEX-NeuroD1-mCherry into the mouse M1 motor cortex and the spinal cord, and then performed a serial immunostaining using both cortical neuronal markers (FoxG1 and Tbr1) and spinal neuronal markers (Tlx3 and Pax2) at 4 wpi (Fig. 4). The majority of NeuroD1-infected cells were converted into neurons in both the brain and the spinal cord at 4 wpi (Fig. 4A, C). Strikingly, when we compared the neuronal subtypes resulting from NeuroD1-mediated conversion in the brain versus the spinal cord side-by-side, a distinct pattern emerged: the converted neurons in the mouse cortex acquired cortical neuron markers such as FoxG1 (66.1 ± 14.3%) and Tbr1 (17.1 ± 1.9%), but not spinal neuron markers such as Tlx3 (0%) or Pax2 (0%) (Fig. 4A, B); in contrast, the converted neurons in the spinal cord acquired spinal neuron markers Tlx3 (46.4 ± 2.2%) and Pax2 (4.2 ± 0.3%), but not cortical neuron markers FoxG1 (0%) or Tbr1 (0.6 ± 0.5%) (Fig. 4C, D). Morphologically, the converted neurons in the brain resembled cortical pyramidal neurons with larger cell bodies (Fig. 4A), while those in the spinal cord resembled dorsal horn interneurons with smaller cell bodies (Fig. 4C). The relative lower percentage of Tbr1+ cells among the converted neurons in the cortex suggest that the newly converted neurons may not be mature enough at 4 wpi and may take longer time to fully acquire their neuronal identity. These distinct differences in the neuronal identity after conversion by the same transcription factor in the brain versus the spinal cord suggest that the glial cell lineage, here cortical lineage versus spinal lineage, as well as the local environment may exert an important influence on the resulting subtypes of converted neurons.

### NeuroD1-converted neurons are physiologically functional

To test the functionality and circuit-integration of NeuroD1-converted neurons, we performed patch-clamp electrophysiological recordings of native and converted neurons on spinal cord slices from mice sacrificed at 8-10 wpi (Fig. 5A). The converted neurons could generate repetitive action potentials (Fig. 5B) and displayed large Na^+^ and K^+^ currents (Fig. 5C). Moreover, we detected robust spontaneous EPSCs from the NeuroD1-converted neurons (Fig. 5D). Quantitatively, we found that the NeuroD1-converted neurons showed similar levels of Na^+^ currents (Fig. 5E) and spontaneous EPSCs to their neighboring native neurons (Fig. 5F). Immunostaining with a series of synaptic markers including SV2 and VGlut1/VGlut2 further confirmed that the NeuroD1-converted neurons were surrounded by numerous synaptic puncta with many of them directly innervating the neuronal soma and dendrites (Fig. 5G, H, cyan and yellow dots). Finally, cFos, an immediate early gene that is typically activated by neuronal activity during functional tasks, was clearly detected in some of the NeuroD1-converted neurons, indicating that they were functionally active in the local spinal cord circuits (Fig. 5I). Altogether, our results demonstrate that NeuroD1 can reprogram reactive astrocytes into functional neurons in the dorsal horn of the injured spinal cord.

**Figure 5:**
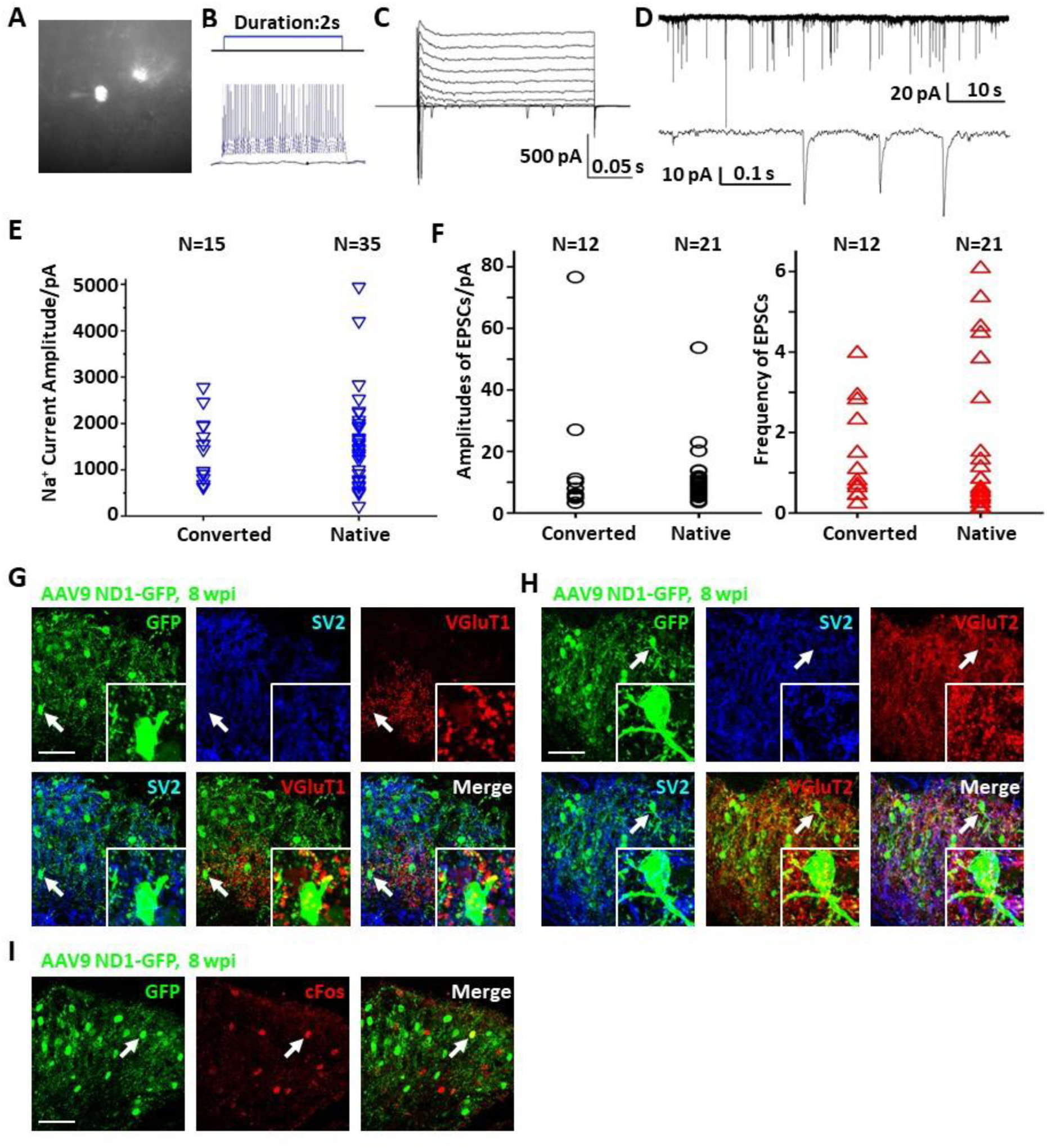
Maturation and functionality of NeuroD1-converted neurons in the spinal cord dorsal horn. (**A**) Fluorescent/transmitted light image of a patch-clamped converted neuron. (**B**) Sample action potentials of a converted neuron. (**C**) Sample Na^+^ and K^+^ currents of a converted neuron. (**D**) Sample EPSCs of a converted neuron. (**E**) Na^+^ current amplitudes for converted and native neurons. (**F**) EPSC amplitudes and frequencies for converted and native neurons. (**G**) Synaptic SV2 and VGluT1 puncta for converted neurons in the dorsal horn 8 wpi after AAV9 ND1-GFP injection. Arrows and inset (4×mag) show an example cell with puncta visible on its soma and processes. Scale bar 50 µm. (**H**) Synaptic SV2 and VGluT2 puncta for converted neurons in the dorsal horn 8 wpi after AAV9 ND1-GFP injection. Arrows and inset (4×mag) show an example cell with puncta visible on its soma and processes. Scale bar 50 µm. (**I**) Integration of converted neurons into local network in the dorsal horn 8 wpi after AAV9 ND1-GFP injection. Activated neurons indicated by cFos staining are only a small subset of all neurons.

### NeuroD1-mediated cell conversion in the contusive SCI model

To move closer towards clinical situations, we evaluated NeuroD1-mediated neuronal conversion in the contusive SCI model. Compared with stab injury, contusive injury creates a much more severe injury environment, which could affect the efficiency of neuronal conversion and the survival of converted neurons. We therefore performed two experiments to test our AAV GFAP∷Cre and Flex-NeuroD1-GFP system after contusive SCI: one short-delay injection to test our treatment as a response to acute injury (Fig. 6) and one long-delay injection to test our treatment as a response to chronic injury (Fig. 7). The advantage of the short-delay experiment is to maximize infection rate by taking advantage of the post-injury proliferation of reactive astrocytes, while the advantage of the long-delay experiment is to maximize the neuronal survival after conversion by allowing injury-induced neuroinflammation to taper down and minimize the secondary effects of the contusion injury. In our short-delay experiment, viral injection was conducted at 10 days post-contusive injury and tissues were collected at 6 weeks post-viral infection (Fig. 6A). Viral injections were performed 1 mm away from the contusion site to avoid the injury core (Fig. 6B). The injury core is apparent after contusion and is characterized by the loss of NeuN+ neuronal cell bodies (Fig. 6C, labeled by *). Viral injection at 10 days post-contusion resulted in many GFP+ cells surrounding the injury core in both control GFP and NeuroD1-GFP groups (Fig. 6C), indicating good infection rate and survival of the AAV-infected cells in the contusive SCI model. On the other hand, the AAV NeuroD1-GFP infected cells showed a dramatic morphological difference from the control GFP group (Fig. 6C). As illustrated in the enlarged images in Fig. 6C, the GFP infected cells in the control group showed typical astrocytic morphology and colocalization with GFAP signal (magenta), but rarely showed any colocalization with the neuronal marker NeuN (red). In contrast, NeuroD1-GFP infected cells were often colocalized with NeuN but rarely colocalized with GFAP (Fig. 6C), indicating successful neuronal conversion. Quantitatively, we counted the total number of converted neurons to be ∼2,600 cells surrounding the lesion core areas (Fig. 6D). The efficiency of NeuroD1-mediated neuronal conversion in the short-delay experiment as measured by NeuN immunoreactivity was ∼55% (Fig. 6E), while the remaining cells were mostly GFAP+ (Fig. 6F). In contrast, the GFP-infected cells were mostly GFAP+ astrocytes and rarely NeuN+ neurons (only 3.9% NeuN+ in GFP group) (Fig. 6E, F).

**Figure 6.**
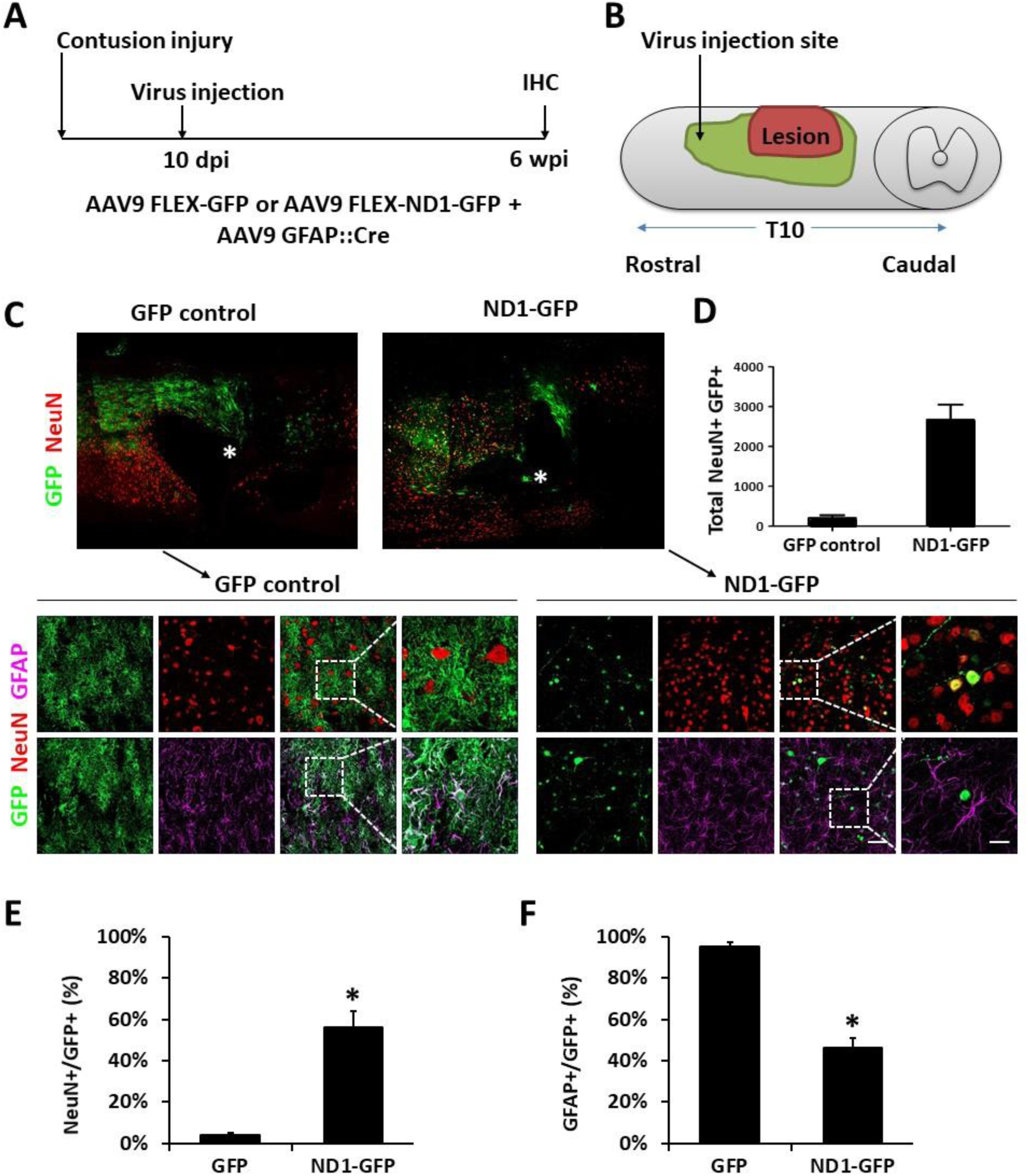
NeuroD1 converts reactive astrocytes into neurons around the injury core with a short delay of viral injection after contusive SCI. (**A**) AAV9 FLEX expressing either GFP reporter alone or NeuroD1-GFP were injected along with AAV9 GFAP∷Cre to target reactive astrocytes at 10 days after a contusive SCI (30 Kdyn force). Spinal cords were analyzed at 6 wpi. (**B**) Experiment paradigm. (**C**) Many infected cells survived around the injury core (indicated by *) and showing distinct cellular morphology between the two groups. Immunostaining of the neuronal markers GFAP and NeuN indicates successful neuronal conversion from reactive astrocytes by ND1-GFP. Scale bars 50 µm at low-mag, 20 µm at high-mag. (**D**) Estimated number of converted neurons per infection (average number of NeuN+ infected cells per horizontal section, calculated from one dorsal, one central, and one ventral section, multiplied by the total number of horizontal sections per sample (n=3 for each group; *, p<0.01). (**E**) NeuN acquisition at 6 wpi (n=3 for each group; *, p<0.01). (**F**) GFAP loss at 6 wpi (n=3 for each group; *, p<0.01).

**Figure 7.**
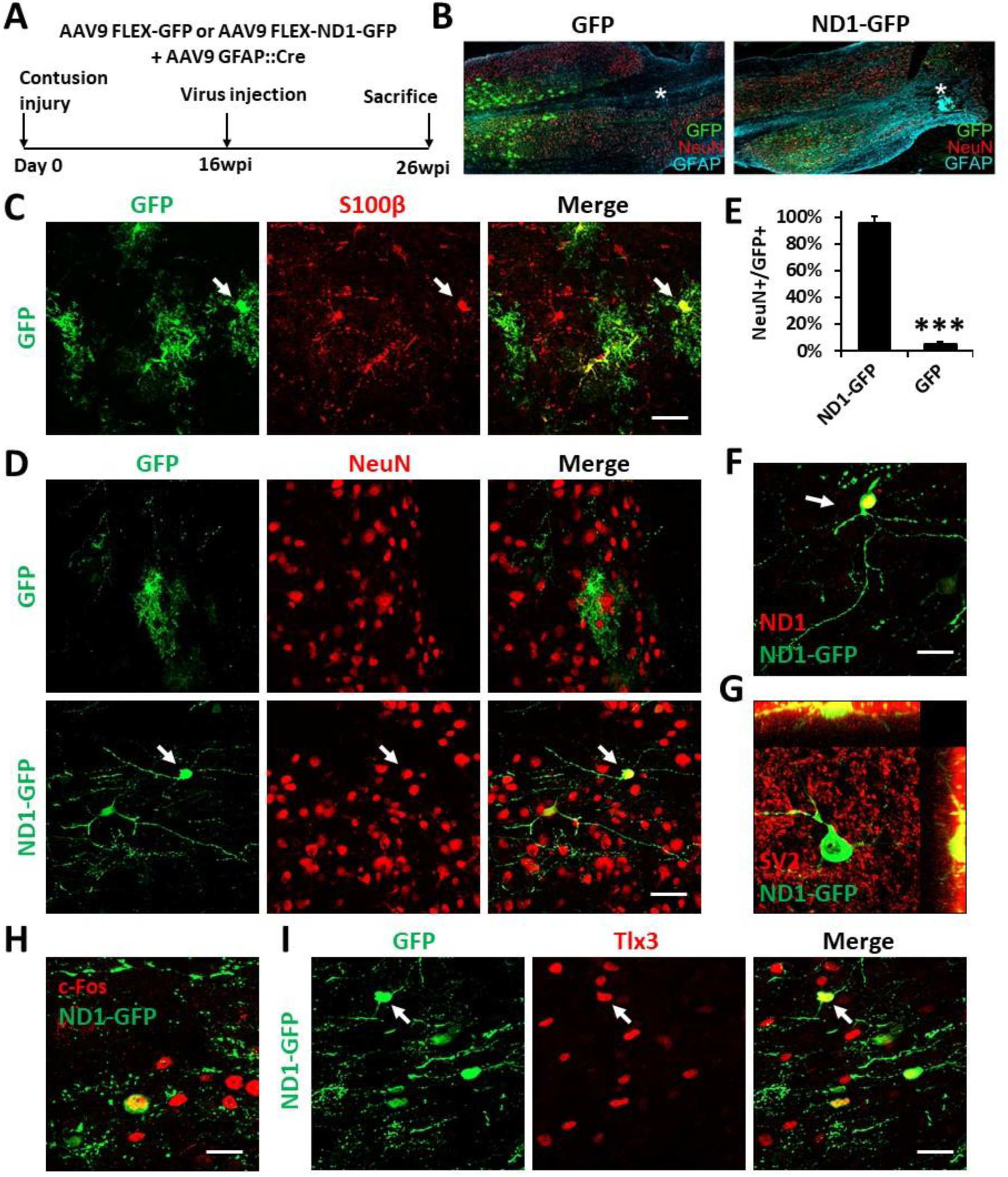
NeuroD1-mediated neuronal conversion with a long delay of viral injection after contusive SCI. (**A**) AAV9 FLEX expressing either GFP reporter alone or NeuroD1-GFP were injected along with AAV9 GFAP∷Cre to target reactive astrocytes at 16 weeks after a contusive SCI (30 Kdyn force). The spinal cords were analyzed at 10 wpi. (**B**) Many infected cells survived around the injury core (indicated by *) and showing distinct cellular morphology between the two groups. (**C**) Co-expression of the astrocyte marker S100b in control GFP+ cells. Scale bar 50 µm. (**D**) Immunostaining of the neuronal marker NeuN indicates successful neuronal conversion from reactive astrocytes by ND1-GFP with high efficiency. Scale bar 50 µm. (**E**) NeuN acquisition at 10 wpi (n=4 for each group; ***, p<0.005). (**F**) Co-expression of ND1 protein in the ND1-GFP+ cells with a typical neuronal morphology. Scale bar 20 µm. (**G**) Co-expression of the mature neuronal marker SV2 in the ND1-GFP+ cells. (**H**) Co-expression of the neuronal activity marker cFos in the ND1-GFP+ cells. Scale bar 20 µm. (**I**) Co-expression of the glutamatergic subtype marker Tlx3 in ND1-GFP+ neurons in the spinal cord dorsal horn. Arrows show an example Tlx3+ cell. Scale bar 20 µm.

In our long-delay experiment, viral injection was conducted at 4 months post-contusive injury, when glial scar has been well formed after contusion, and tissues were collected at 10 weeks post-viral infection (Fig. 7A). Fig. 7B illustrates the overall morphology of the spinal cord with immunostaining of GFP, NeuN, and GFAP. As in the short-delay experiment, the lesion core (labeled by *) also lacked NeuN+ neurons. In the control AAV GFP alone group, the viral infected cells were mainly S100b+ astrocytes (Fig. 7C), but rarely showed any NeuN+ signal (Fig. 7D, top row). In contrast, the majority of NeuroD1-GFP infected cells were converted into NeuN+ neurons (Fig. 7D, bottom row; quantified in Fig. 7E). The NeuroD1-mediated conversion efficiency reached >95% (Fig. 7E). Immunostaining confirmed the NeuroD1 overexpression in the NeuroD1-GFP infected cells (Fig. 7F). Furthermore, NeuroD1-converted neurons at 10 wpi were surrounded by many synaptic puncta (SV2) with some of them directly innervating the soma and dendrites (Fig. 7G, yellow dots). We also identified c-Fos+ cells among NeuroD1-converted neurons (Fig. 7H), indicating that they were able to integrate into the local spinal cord functional circuitry. Lastly, some of the NeuroD1-converted neurons in the contusive SCI model at 10 wpi showed glutamatergic subtype through expression of Tlx3 in the dorsal horn (Fig. 7I), consistent with our stab injury model. Altogether, these results indicate that NeuroD1 overexpression can reprogram reactive astrocytes into functional neurons after contusive SCI under both acute and chronic treatment conditions, with higher conversion efficiency achieved after glial scar formation. This clinically relevant model can be used in future studies to further test functional improvement after SCI using *in vivo* cell conversion technology.

## Discussion

In the current study, we have demonstrated in different SCI models that overexpression of NeuroD1 in the reactive astrocytes can convert them into Tlx3-positive glutamatergic neurons in the dorsal horn of the injured spinal cord. Other ongoing studies are also testing different transcription factors to reprogram spinal astrocytes into GABAergic neurons or motor neurons. Using the AAV Cre-Flex system, we efficiently reprogrammed reactive astrocytes around the injury site in the dorsal horn of the spinal cord into functional neurons that integrate into the local synaptic network. Importantly, we also observed highly efficient NeuroD1-mediated neuronal conversion in the contusive SCI model making this technique a feasible intervention for regenerating functional new neurons in the grey matter to treat SCI. Consistent with our *in vivo* cell conversion work in the brain (Chen et al., 2018; Guo et al., 2014), the NeuroD1-converted neurons in the dorsal horn of the injured spinal cord were also glutamatergic, but Tlx3+ spinal neurons instead of Tbr1+ cortical neurons. The fact that the same neural transcription factor NeuroD1 can convert cortical astrocytes into cortical neurons and spinal astrocytes into spinal neurons may hold the key to a region-flexible neural repair by using internal glial cells for neuroregeneration. In contrast, transplantation of external cells does not have this advantage of intrinsic cell lineage.

### AAV gene delivery system for neuronal conversion

We have accomplished successful NeuroD1-mediated neuronal conversion with retroviral vectors in both this and previous studies (Guo et al., 2014). However, since retrovirus particles are relatively large, limiting the viral titer, and since retrovirus only infects proliferating cells, the viral injection timing is confined to a narrow time window after the injury during which glial cell proliferation is increased. In contrast, because AAV infects both proliferating and quiescent cells, the AAV injection depends less on timing and can be performed in both acute and chronic conditions. Additionally, AAV has passed clinical trials and elicits little immune response when applied *in vivo* (High and Roncarolo, 2019; Zaiss et al., 2002). AAV particles are small and can be prepared at very high titer, therefore high dosage of conversion factors can be achieved when multiple AAV particles infect a single cell. Consistent with this, we observed a slower neuronal conversion when a lower titer of AAV-NeuroD1 was injected in the injured spinal cord (unpublished observation). Since certain serotypes of AAV can cross the blood-brain-barrier (BBB), intravenous delivery of conversion factors has make it possible to reach a broader area of the CNS (Chan et al., 2017; Foust et al., 2009). Following our earlier report of efficient conversion of reactive astrocytes into functional neurons by NeuroD1 through retroviral infection (Guo et al., 2014), intravenous injection of AAV9 expressing NeuroD1 has been reported to infect a small but significant number of resting astrocytes in the striatum and convert them into neurons (Brulet et al., 2017). On the other hand, our intracranial injection of AAV9 expressing NeuroD1 in stab injury model (Zhang et al., 2018) and ischemic injury model (Chen et al., 2018) have both resulted in high conversion efficiency, suggesting that reactive astrocytes are more likely converted into neurons than the resting astrocytes. Consistent with this hypothesis, our results in this study also support that reactive astrocytes in injured spinal cord can be effectively converted into neurons with ∼95% efficiency, regardless of retrovirus or AAV delivery system.

### Distinct functions of NeuroD1 during neuronal conversion

Neuronal conversion can be achieved by several neurogenic transcription factors (Li and Chen, 2016). Besides NeuroD1, Sox2, Ngn2, and Ascl1 have all been reported to convert glial cells into neurons (Gascon et al., 2016; Grande et al., 2013; Guo et al., 2014; Heinrich et al., 2014; Liu et al., 2015; Niu et al., 2013; Su et al., 2014). Sox2 is expressed in neural progenitors and functions to maintain progenitor identity (Bani-Yaghoub et al., 2006; Bylund et al., 2003; Graham et al., 2003). Therefore, it is not surprising that Sox2-mediated neuronal conversion has to go through a proliferation stage as shown by incorporation of BrdU and the expression of Ki67 (Su et al., 2014). In contrast, NeuroD1 is a neuronal differentiation transcription factor that instructs terminal differentiation of neuroprogenitors into neurons during early neural development (Gao et al., 2009; Miyata et al., 1999; Morrow et al., 1999). This might partially explain why NeuroD1 can achieve high neuronal conversion efficiency (>90%) comparing to ∼6% in the case of Sox2 (Su et al., 2014). An earlier report also showed that Ngn2-expressing retrovirus was able to promote neurogenesis in the injured spinal cord, but the number of newly generated neurons greatly decreases over time even when combined with growth factor treatment (Ohori et al., 2006). When combined with Bcl2, an anti-apoptotic gene, Ngn2-mediated neuronal conversion acquires a much higher efficiency, although the authors claim that Bcl2 plays a role independent of apoptotic pathways (Gascon et al., 2016). In sharp contrast, we rarely observe apoptotic cells during and after NeuroD1-mediated conversion as determined by TUNEL assay. The difference of cell survival in converted neurons between different transcription factors may be explained by the fact that NeuroD1 is not only a conversion factor but also a survival factor. During development, NeuroD1 is required for survival of a variety of neuronal subtypes in the developing and adult CNS (Gao et al., 2009; Miyata et al., 1999; Morrow et al., 1999). This dual role of NeuroD1 during neuronal conversion and neuronal survival may explain its higher conversion efficiency over Sox2 and Ngn2. Ascl1 has also been reported to induce high efficiency of *in vivo* astrocyte-to-neuron conversion in the midbrain (Liu et al., 2015), suggesting that Ascl1 might share certain common properties with NeuroD1. Since long-term survival of converted neurons is essential to their integration into local neuronal circuitry in order to have a role in functional repair, future studies on clinical translation must pay much attention to the total number of newly generated neurons that can survive for years to have effective therapies.

### Environmental cues to impact neuronal conversion in addition to intrinsic factors

Transcription factor-mediated *in vivo* neuronal reprogramming illustrates intrinsic power to convert reactive astrocytes into neurons. However, environmental cues also play a role in the success of neuronal conversion (Heinrich et al., 2014) as well as the phenotype of converted neurons (Grande et al., 2013). In this study, we found that NeuroD1-converted neurons in the injured mouse cortex were Tbr1+ cortical neurons, but in the injured spinal cord the NeuroD1-converted neurons were Tlx3+ spinal neurons. Therefore, the local environment, together with astroglial lineage, may be essential to functional integration of converted neurons into the local neuronal circuitry as they mature. Together, a complete neuronal conversion would need both intrinsic factors (transcription factors and glial lineage factors) and extrinsic factors (local cues) to solidify the identity of converted neurons.

Even within the spinal cord, local environment can be drastically different between the grey matter versus the white matter, with neuronal soma confined to the grey matter and neuronal axons occupying the white matter. In our experiments using AAV, we rarely observed converted neurons in the white matter; using retrovirus, we observed some converted neurons in the white matter at early time points, but they rarely survived to 6 wpi. This has been similarly observed in cell conversion studies in the white matter (corpus callosum) of mouse brains (Liu et al., 2020). Reasons for the lack of conversion in the white matter in this study could include our targeted injection technique which delivers virus precisely into the dorsal horn of the grey matter or the lack of appropriate viral receptors in the white matter. White matter may also lack sufficient trophic factors for the survival of newly generated neurons. On the other hand, Sox2-mediated neuronal conversion can result in many newborn neurons located in the white matter of the spinal cord, particularly in p21 knockout mice (Wang et al., 2016). An interesting feature of these neurons is that they appear as clusters, which, by providing trophic factors to each other, could be the reason they survive. It is also possible that the local environment in the white matter of p21 knockout mice has been altered during Sox2-mediated neuronal conversion in combination with BDNF and noggin. Further studies will be required to determine the differential effects of not only grey matter versus white matter but also dorsal horn versus ventral horn on neuronal conversion.

### Generation of neuronal subtypes via NeuroD1-mediated conversion

We demonstrate here that NeuroD1 converts reactive astrocytes into primarily Tlx3-positive glutamatergic neurons in the dorsal horn of the injured spinal cord. Interestingly, Sox2-converted neurons in the injured spinal cord are also mainly glutamatergic (Wang et al., 2016), raising the possibility that glutamatergic neurons might be a default subtype of converted neurons. On the other hand, one cannot ignore the small yet significant component of GABAergic neurons (∼10-15%) generated by NeuroD1 in the dorsal horn of spinal cord, which we found to be roughly equal to the population in our non-injured control and might still play a meaningful role in the local circuitry. Although NeuroD1 overexpression has been shown to inhibit GABAergic neuronal differentiation by suppressing Ascl1 (Mash1) (Roybon et al., 2010), we have consistently found that NeuroD1 can convert at least 10% of glial cells into GABAergic neurons, suggesting that astrocyte-to-neuron conversion is distinct from stem cell differentiation in terms of potential mechanisms. Of course, in order to generate more GABAergic neurons, combinations with other transcription factors may be necessary; this is indeed an ongoing study in this lab.

## Conclusion

In summary, our study demonstrates that NeuroD1 can convert reactive astrocytes into neurons in the injured spinal cord, and that the AAV gene delivery system does so with high conversion efficiency. Furthermore, the AAV-NeuroD1 converted neurons can functionally mature and integrate into the local spinal cord network. We also show that AAV NeuroD1-converted neurons in the spinal cord dorsal horn mainly acquire a Tlx3+ glutamatergic neuronal subtype. However, we believe that a diversity of neuronal subtypes including GABAergic neurons and ventral horn motor neurons are needed to replenish lost neurons after SCI in order to reconstruct damaged neuronal circuitry and restore spinal cord functions. With a therapeutic aim in mind, our next challenge is to optimize this *in vivo* neuronal conversion technology in terms of combinations with other factors, viral injection dosage and timing, and ultimately its ability to lead to beneficial effect and functional improvement in various SCI models.

## Acknowledgement

This work was mainly supported by Charles H. Smith Endowment Fund for Brain Repair (G.C.) and Verne M. Willaman Endowment Fund from the Pennsylvania State University (G.C.), as well as a grant from National Institutes of Health R21NS104394 (H.L.).

## Supplemental figures

**Supplemental Figure 1.**
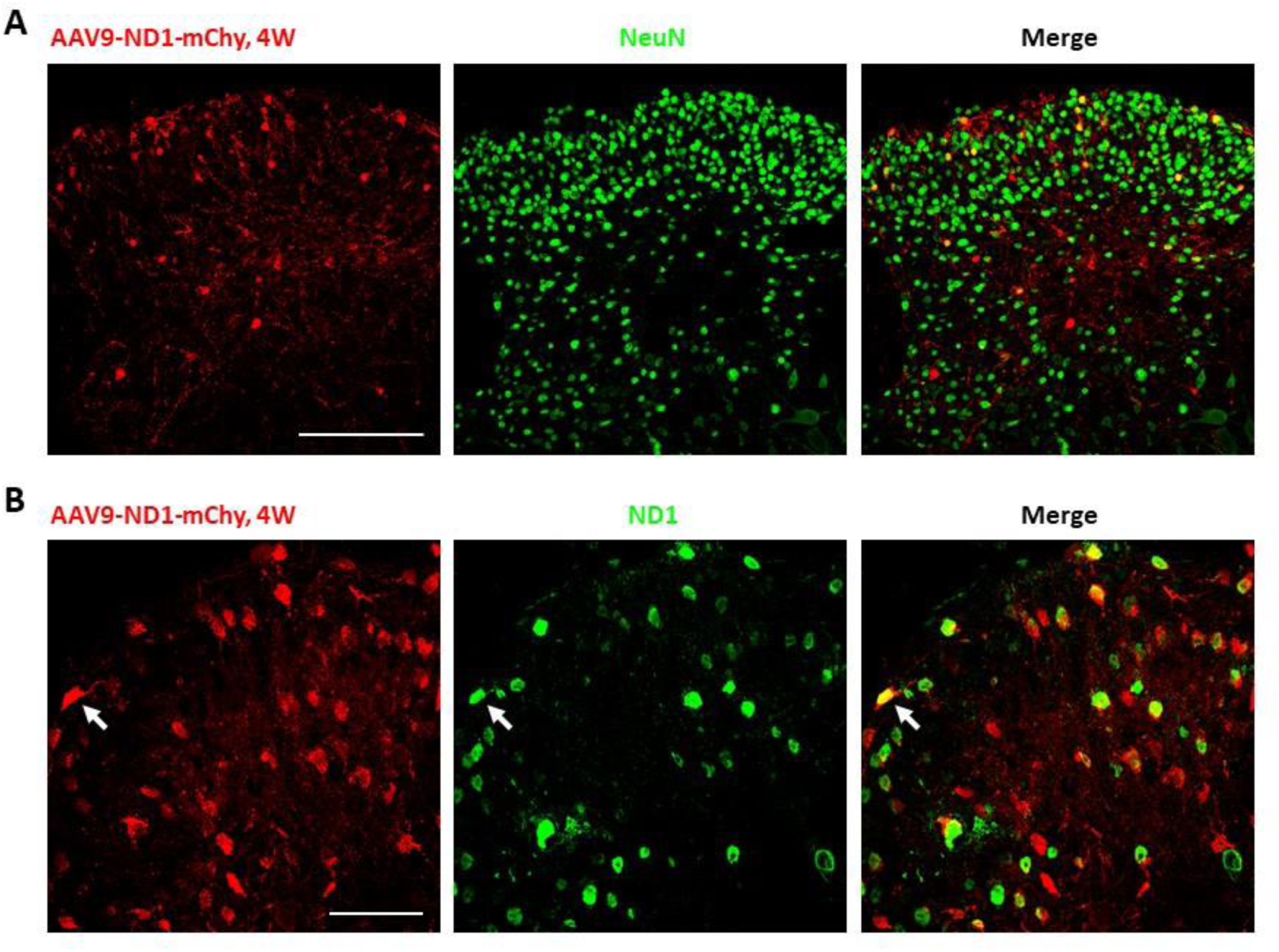
Infected cells by AAV9-ND1-mChy overexpress ND1 protein in the injured spinal cord. (**A**) Immunostaining analysis confirmed that the infected cells by AAV9 ND1-mCh expressed the neuronal marker NeuN indicating neuronal conversion in the dorsal horn of injured spinal cord at 4 wpi. Scale bar 200 µm. (**B**) Infected cells overexpressed ND1 protein at 4 wpi. Scale bar 50 µm.

**Supplemental Figure 2.**
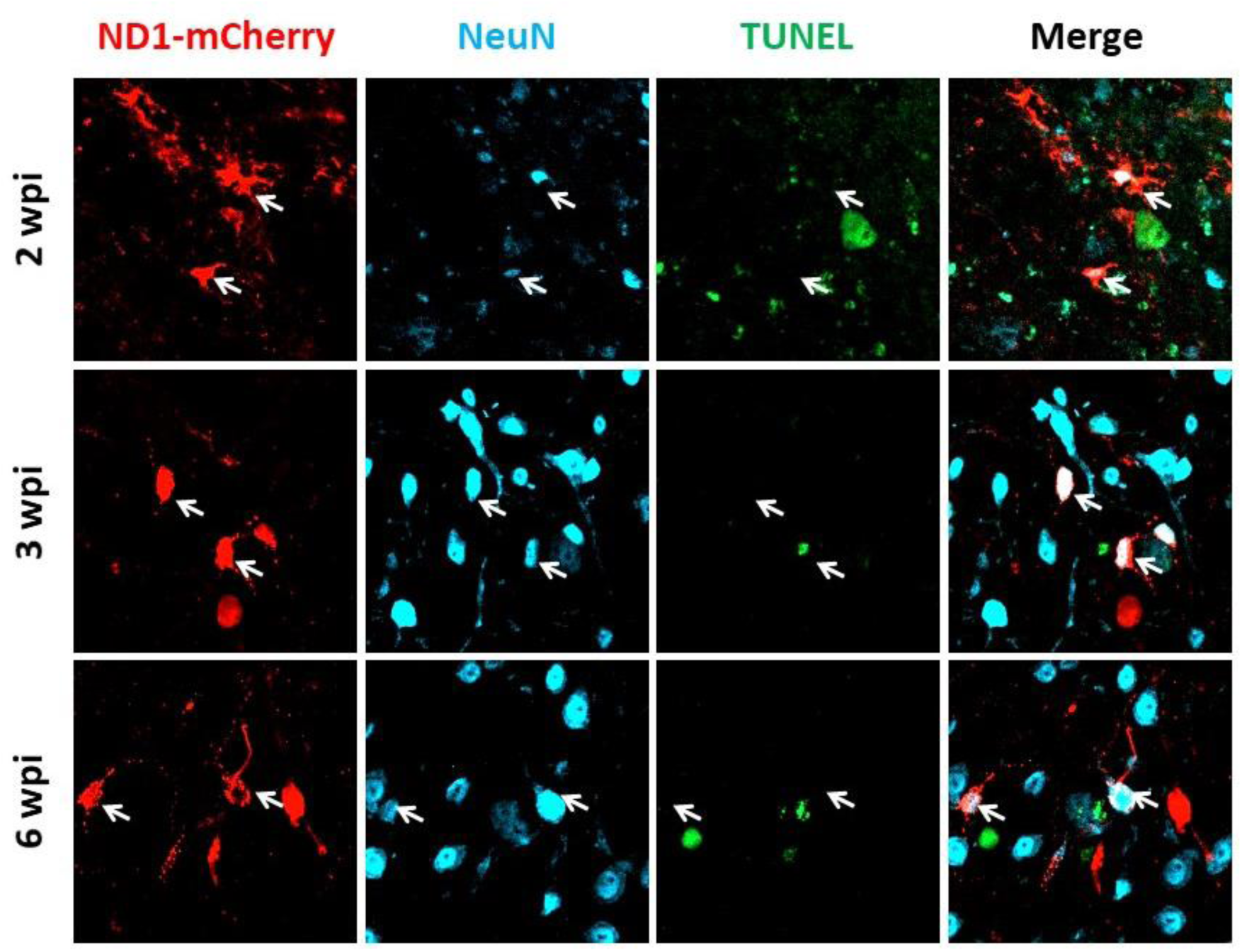
NeuroD1-mediated neuronal conversion does not involve severe apoptosis. TUNEL assays were performed to detect apoptotic cells at different stages of neuronal conversion by AAV9 ND1-mCh in the injured spinal cord. Arrows show infected cells that are NeuN+ but TUNEL−.

**Supplemental Figure 3.**
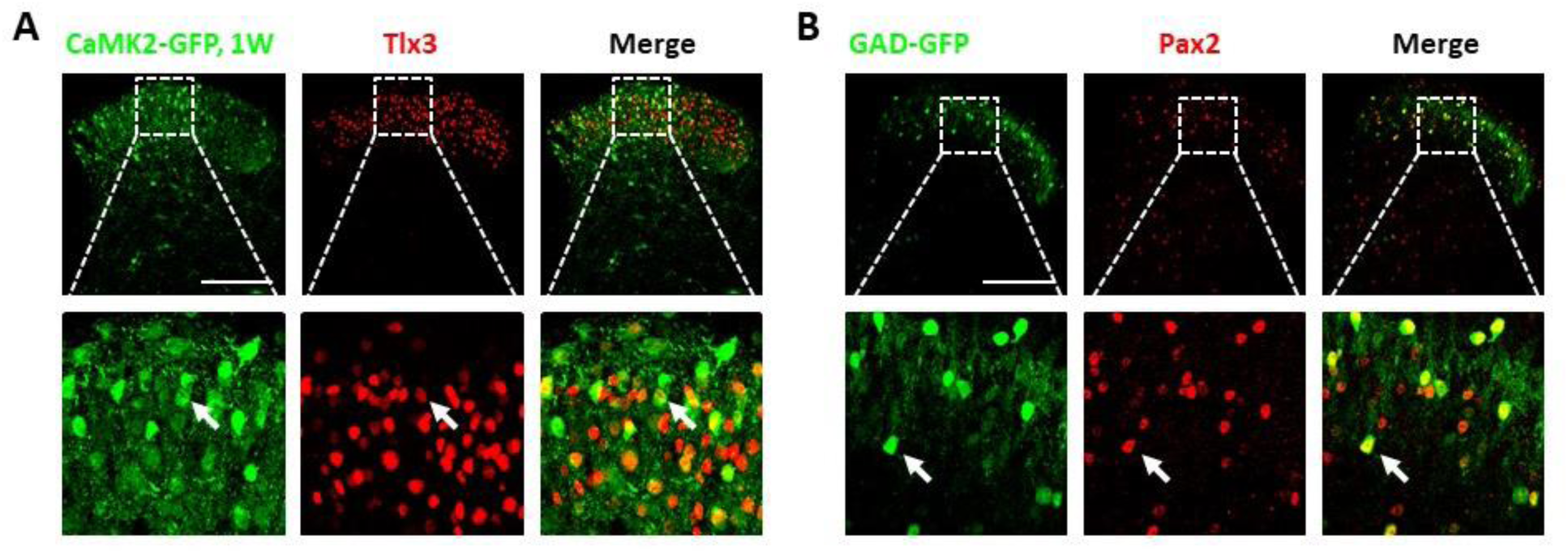
CaMK2-GFP virus and GAD-GFP mice can be used to confirm neuronal subtype. (**A**) CaMK2-GFP (from co-injected AAV9 virus) co-stains with Tlx3 while (**B**) GAD-GFP (from GAD-GFP transgenic mouse) co-stains with Pax2, indicating that these markers can be used to confirm glutamatergic and GABAergic subtypes in the dorsal horn. Scale bar 200 µm. Dotted boxes at 4×mag below.

## Notes

**Conflict of Interest:** G.C. is a co-founder of NeuExcell Therapeutics.

#### Summary of Updates

Author list

